# Insect attraction to the six major types of traditional-style, residential light bulbs and implications for insect survival and light pollution

**DOI:** 10.1101/2021.05.06.442978

**Authors:** Michael J. Justice, Teresa C. Justice

**Affiliations:** Unaffiliated

**Keywords:** Incandescent, CFL, Halogen, Bug Light, Light Pollution, Urban Ecology

## Abstract

Artificial light at night can affect the behavior and survival of the arthropods attracted to it. Most light pollution research focuses on high-wattage street lamps, but lower-wattage lamps used to illuminate porches, paths, façades, and backyards vastly outnumber street lamps. Thus, residential consumers could potentially have an enormous influence on artificial light ecologies by their choice of lamp. This study compared insect attraction to the six major types of traditional-style, residential light bulb: incandescent, CFL, halogen, warm color temperature LED, cool color temperature LED, and the yellow “bug” lights marketed as reducing insect attraction. The bulbs were alternately used in a baffle-funnel light trap from early spring through late fall, and capture rates were determined for the orders of insects. Incandescent bulbs produced the highest trap captures; the warm color temperature LED bulb produced the fewest, even fewer than the yellow “bug” light. The yellow “bug” light attracted more Dermaptera (Leach) than the other bulbs. The data support a recommendation of LED bulbs, especially those with a warm color temperature, to minimize the effects of night lighting on insect behavior and mortality. Further, the use of yellow “bug” lights, in contrast to their marketing, could attract earwigs and other minor pests.

---

Insects evolved in an ecology lit mostly by the sun and moon. “Artificial light ecologies” have resulted from the widespread use of artificial light at night, which is often referred to as “ecological light pollution” (Rich & Longcore 2006). Insects are affected by light pollution in many ways. Some lamps operate at temperatures hot enough to desiccate, injure, or kill insects that approach or alight on them. Time spent around lamps may reduce the time spent in normal behaviors; among insects, changes in hatching, feeding, dispersal, pupation, eclosion, mate searching, mating, and oviposition have been documented or suspected (Frank 1988, 2006; Perkin et al. 2013; van Geffen et al. 2014). The effects of light pollution on individual insects could extend to populations (Frank 2006) and communities (Buchanan 2006; Perry & Fisher 2006; Rydell 2006; Zhang et al. 2010; Davies et al. 2012). Such effects can weigh heavily in ecosystems, agriculture, and disease transmission, so it is important to explore ways in which they can be mitigated.

Many types of lamp are available to consumers to illuminate small areas around homes such as porches, paths, façades, and yards. As the luminaires are often already installed, many lamps share the traditional A19 shape (commonly referred to as a “bulb”) and the E26 screw base. Understanding the nature of insect attraction to such lamps is important because even a small difference in insect attraction may become profound when multiplied by the vast numbers of such lamps that are in regular use. A consumer’s choice may thus be a small contribution to a larger ecological problem. Assuming a light bulb purchaser needs a certain level of brightness (which is indicated by lumens on the packaging), the main choices then are “color temperature” and the mechanism by which light is produced. Color temperature uses the Kelvin temperature of a blackbody radiator as a reference. Bulbs with “Warm” color temperatures (approximating a blackbody radiator at ~3000K) are more red. The bluer “Cool” color temperatures (~5000K) are more susceptible to atmospheric scattering and thus worse for light pollution (Falchi et al. 2011).

Incandescent bulbs have a warm color temperature. Due to their extreme energy inefficiency, they are being phased out of many markets. Modern replacements have better energy efficiency and offer a wider range of color temperatures. Halogen lamps are modified incandescent lamps that also have high operating temperatures and offer only slightly better energy efficiency. More recent technologies create short-wavelength/UV radiation which causes a mix of powdered phosphors coating the bulb to fluoresce. Different color temperatures are produced by altering the ratio of the phosphors. The short-wavelength radiation is created by ionizing mercury in compact fluorescent lights (CFLs) and by solid-state electronics in light-emitting diodes (LEDs).

In addition to lamps that produce some variant of white light, lamps that are yellow in appearance are sold as “bug lights.” These are marketed as either reducing insect attraction or actively repelling insects. This is presumably based on either 1) insects’ visual systems being in general more UV-blue sensitive and less yellow-red sensitive, or 2) the lamps’ similarity in appearance to high-wattage, sodium metal-halide street lights, which have relatively low levels of insect attraction (Rich & Longcore 2006). These presumptions may be reasonable, but insect attraction to lower-wattage yellow bulbs that use different mechanisms of light production has not been systematically quantified.

Studies specifically examining the low-wattage light bulbs designed for residential areas are scarce (Truxa & Fielder 2012; van Geffen et al. 2014). Much of the light pollution research has focused on street lighting, especially comparing sodium and mercury vapor lamps to other high-intensity lamps (e.g. Eisenbeis 2006; Eisenbeis & Eick 2011; Perkin et al. 2013). Studies directed at residential lighting generally involve short-wavelength/UV “blacklights” because of their use in traps designed to reduce nuisance insects (e.g. Hienton 1974; Nabli et al. 1999). Justice & Justice (2016), using residential-type light bulbs, found that LED lamps produced the lowest insect capture rates. This study replicates and extends these results by using a much longer collecting period and directly comparing five types of “white” light bulbs to a yellow “bug” light.

## Materials and Methods

This study compared six light bulbs that can be readily purchased at retail stores in the United States (Table 1). All were labelled as 790-800 lumens brightness and thus from a consumer perspective at the point of purchase would be considered functionally equivalent for illumination purposes.

**Table 1.** Lamps used in the present study. All had the A19 “bulb” shape and E26 screw base. “Cool” and “Warm” refer to color temperature (CT). For comparison, the color temperature of moonlight is approximately 4000 K (Johnsen 2012). CTs were provided by the manufacturer for all bulbs except the yellow bug light.

| Light Bulb Type | Manufacturer | Product Number | Watts | Lumens | CT (K) |
| --- | --- | --- | --- | --- | --- |
| Incandescent | Sylvania | 60A/CL/DL/RP | 60 | 790 | 2850 |
| CFL | Utilitech | 0296865 | 15 | 800 | 3500 |
| Halogen | Sylvania | 43A/HAL/CL/2 | 43 | 800 | 2900 |
| Warm LED | Utilitech | 0383017 | 12 | 800 | 3000 |
| Cool LED | Utilitech | 0383016 | 12 | 800 | 5000 |
| Yellow “Bug” CFL | Sylvania | CF13EL/MINI/BUG/BL/ZJ | 13 | 800 | N/A |

The emission spectrum of each bulb is presented in Figure 1. To obtain the spectra, a luminaire was securely mounted so that the bulb was positioned near the input slit of a Jarrell-Ash 82-410 monochromator. A diffraction grating (1180 grooves/mm) separated the light into its component wavelengths and directed individual wavelengths through a 150 μm output slit. A silicon PIN photodiode detected the intensity at the output slit over the wavelength range 300-750 nm, which covers the known visual range of animal eyes (Land & Nilsson 2012). Amplified voltage was digitized and recorded using a PASCO 750 interface and Data Studio software. A sampling rate of 20 Hz scanned 3.36 nm/sec and collected 5.95 data points per nm. Results were plotted with Microsoft Excel. This system was repeatedly calibrated by generating a spectrum with an OSRAM mercury lamp and comparing it with published spectra for mercury.

**Fig. 1.**
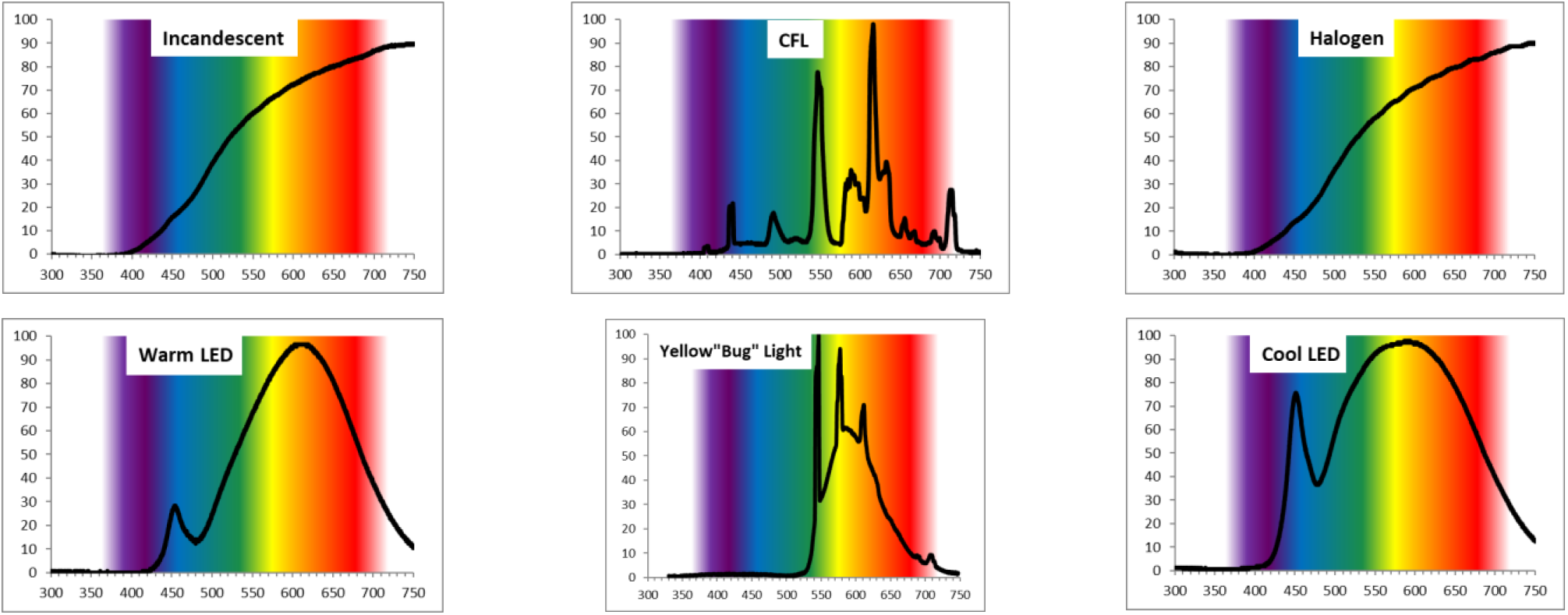
Spectra of the lamps used in the present study. *x*-axes are wavelength in nm, *y*-axes are percent maximum (based on the gain on the photodiode amplifier). It should be noted that none of the bulbs produces much EM radiation below 400 nm, which when present may be an important attractant for insects (Barghini and de Medeiros 2012).

A baffle/funnel gravity-type light trap was created by modifying a Universal Black Light Trap from BioQuip (product #2851A). An aluminium dome holding a lamp socket sat atop the insect baffle, and a jar with 91% isopropyl alcohol was placed at the base of the funnel. A CFL-ready, dusk-to-dawn photoelectric switch regulated 120V AC power to the socket. To protect the trap from rain, it was placed under a carport canopy.

The study was conducted in the town center of Appomattox (37°21’13.5”N, 78°49’32”W), Virginia, USA. As there are only a few very small bodies of water near the study site, aquatic insects were not typically collected. Past research has shown that other lamps in the area of a light trap can affect capture rates by increasing overall brightness or attracting arthropods away from the trap (Rydell 2006), although the diameter of the affected area around each lamp is disputed (Frank 1988, 2006). During the course of this study, no other lamps were ever lit within 50 m of the trap. The dominant light around the study site came from sodium vapor street lamps.

Trapping took place every night between 26 Apr and 28 Sep 2014 with one of the six bulbs assigned *a priori* to each night. Bulbs were initially scheduled so that 1) bulb types were similarly distributed across the study period, 2) no bulb was ever run on consecutive nights, and 3) the moon’s irradiance was equalized across bulb types (especially, each bulb was used at least once in the ±3 days around the new and full moons). For the latter, the percent of the moon’s visible disk that was illuminated at midnight was obtained from the U.S. Naval Observatory’s Astronomical Applications Department. Bright moonlight often reduces captures at light traps (Yela & Holyoak 1997; Eisenbeis 2006), although whether moonlight reduces insect activity, reduces catchment areas, or improves orientation is still unresolved (Nowinszky 2004). Because the irradiance of the illuminated portion of the moon varies with phase angle (Johnsen 2012), and a waxing moon is brighter than a waning moon at the same phase angle (Lane & Irvine 1973; Lawrence et al. 2003), the bulb schedule attempted to equalize percentage of the moon’s disk illuminated in the waxing and waning phases.

As the study progressed, several meteorological variables were monitored using weather data obtained from the Appomattox Court House weather station (37°21’21”N, 78°49’53”W; 0.57 km from trapping site) via the Past Weather service at weather.com. The bulb schedule was occasionally altered to try to correct any treatment differences that developed (without, however, altering the scheduling parameters #1-3 mentioned above). In the end, the 156 nights of trapping were divided as follows: incandescent bulb n = 25 nights, CFL n = 27, halogen n = 25, warm LED n = 26, cool LED n = 25, and yellow n = 28. Treatment differences in weather variables were reduced but not fully resolved; these were addressed statistically (see below).

The captured insects were identified to order using the key in Borror & White (1970) and a dissecting microscope with 7-40X magnification. All insects were counted except mites (Acari), thrips (Thysanoptera, Haliday), and springtails (Collembola, Lubbock), which were highly variable, occasionally innumerable, and, in the case of mites, may have simply arrived on a captured host. For each night, the counts were divided by duration of darkness (as provided by the U.S. Naval Observatory’s Astronomical Applications Department), which expressed them as capture rates and corrected for changing durations of darkness through the summer.

Replicates were not used because, on a given night, setting out more lamps at sufficient distances away to ensure attraction radii do not overlap would have placed the traps in such different habitats that unwanted variability may have been introduced. Between treatment conditions, the percentage of hourly weather reports of rain, clouds, and fog were the most unequal, variable, and concerning (Table 2).

**Table 2.**
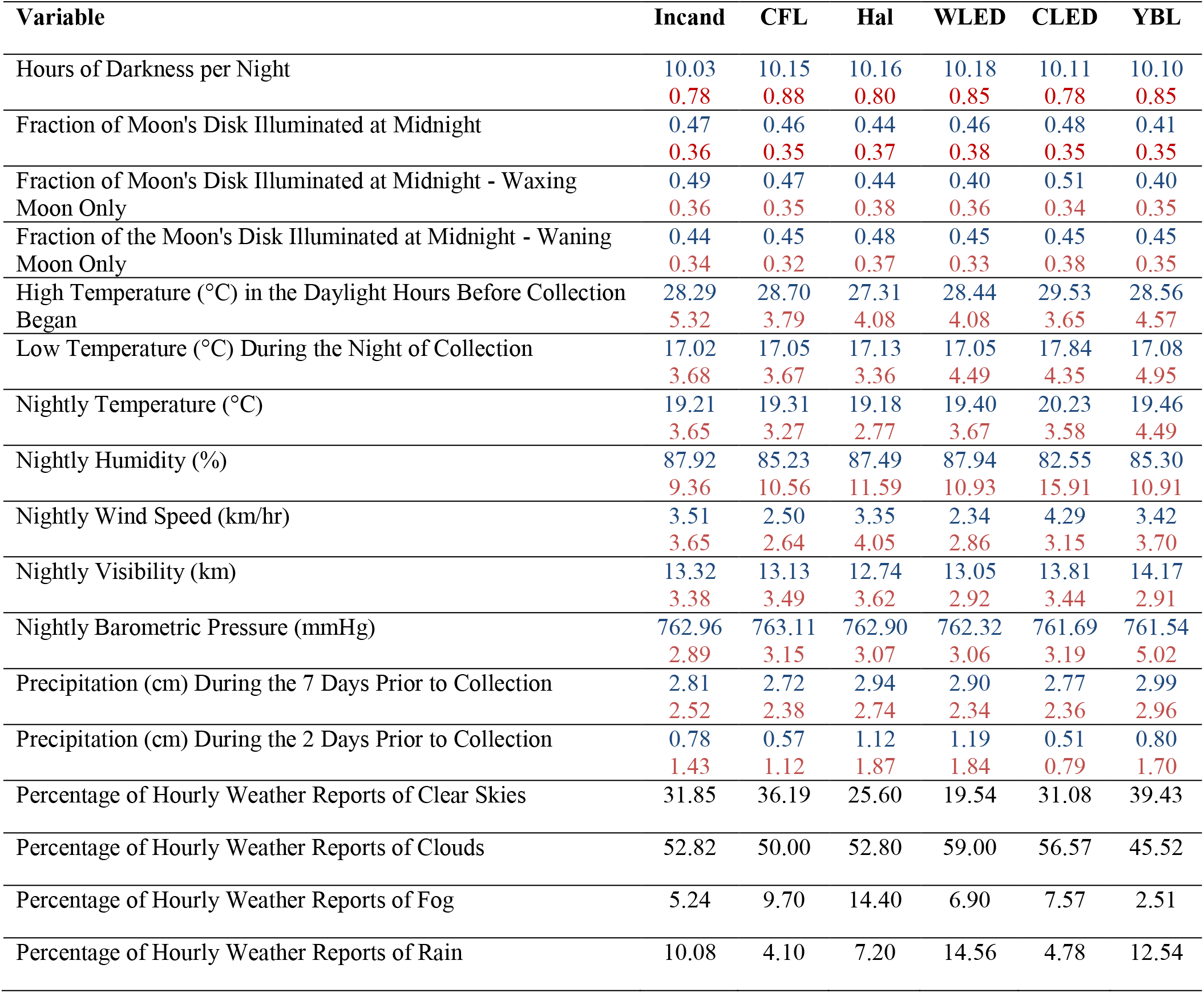
Astronomical and meteorological variables arranged by lamp treatment condition. Incand = Incandescent, CFL = Compact Fluorescent Light, Hal = Halogen, WLED = Warm color temperature LED, CLED = Cool color temperature LED, YBL = Yellow “Bug” Light. “Nightly” indicates that statistics are calculated using the averaged hourly reports from the weather station during the hours of darkness (similar to the method used by Perkin et al. 2013). Where two numbers are present, they are Mean over Standard Deviation.

To address this, weather during collection time was used as a covariate in the analyses (entered as the nightly percentage of hourly weather reports of clear skies; referred to below as the “weather covariate”). The weather variable with the highest correlation with the capture rates was the high temperature on the day collection began, so this was also explored as a covariate (the “temperature covariate”). However, these two weather variables were not used as covariates simultaneously to avoid collinearity. Parametric ANCOVAs and post-hoc tests were used to test for mean differences if samples or their transforms were normally distributed and homoscedastic; otherwise, nonparametric rank-based equivalents were used. Statistical analyses were performed using either Microsoft Excel or computational methods described by Sokal & Rohlf (1981), Rohlf & Sokal (1981), Siegel (1956), and Kirk (1995).

## Results

A total of 8887 insects and spiders were captured, an average of 57 per night (median = 46, *s* = 39.9, minimum = 2, maximum = 246). Average nightly captures per hour of darkness = 5.7 (median = 4.7, *s* = 4.2, minimum = 0.2, maximum = 26.2). The rate of insects and spiders captured was different between the lamp treatment groups (Figure 2).

**Fig. 2.**
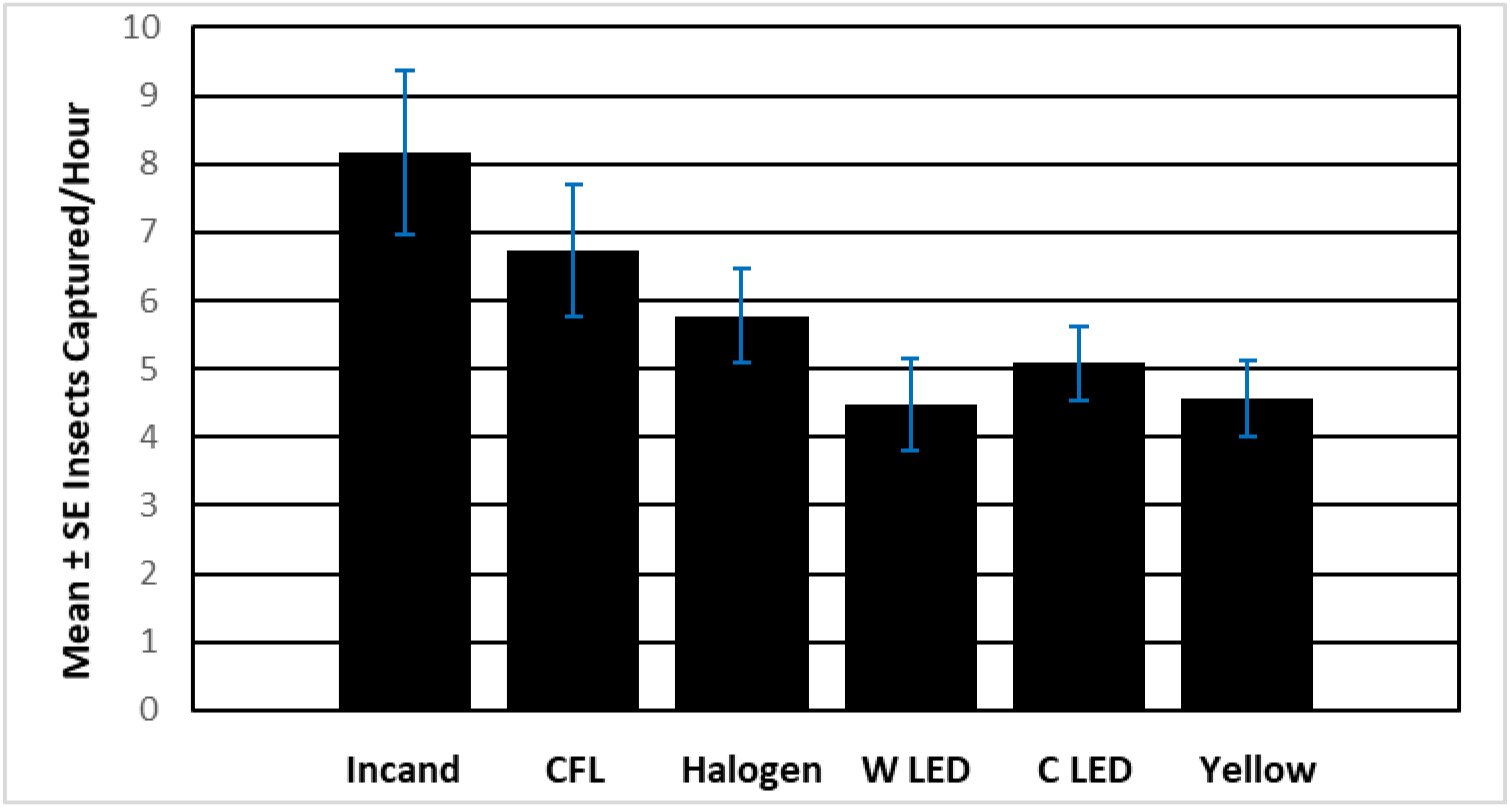
Nightly total captures by bulb type. For each night entered into the average, the total number of captures was divided by the hours of darkness that evening.

The mean ± SE nightly captures/hr was 8.18 ± 1.20 for Incandescent, 6.73 ± 0.96 for CFL, 5.78 ± 0.70 for Halogen, 4.47 ± 0.67 for Warm LED, 5.08 ± 0.54 for Cool LED, and 4.56 ± 0.56 for Yellow. Sample distributions were heteroscedastic (Brown-Forsythe test, *F* = 2.52; df = 5,150; *P* = 0.03), but log-transformed data were homoscedastic (Brown-Forsythe test, *F* = 0.61; df = 5,150; *P* = 0.69) and normally distributed (Kolmogorov-Smirnov *D* statistics, *P* values on all six tests > 0.20). A parametric ANCOVA with the weather covariate yielded *F* = 2.1995; df = 5,149; *P* = 0.0573. With the temperature covariate, *F* = 6.88; df = 5,149; *P* < 0.001. Pairwise comparisons were performed on the log transforms using the Bryant-Paulson generalization of Tukey’s HSD test with the temperature covariate as a random effect and an experimentwise α = 0.05. Three comparisons were significant: Incandescent > Warm LED, Incandescent > Yellow, and CFL > Warm LED.

The rate of Diptera (Börner) captured was different between the lamp treatment groups (total n = 2325 captured). The mean ± SE nightly captures/hr was 2.49 ± 0.39 for Incandescent, 1.79 ± 0.27 for CFL, 1.49 ± 0.26 for Halogen, 1.13 ± 0.21 for Warm LED, 1.33 ± 0.22 for Cool LED, and 0.75 ± 0.12 for Yellow. Sample distributions were heteroscedastic (Brown-Forsythe test, *F* = 4.05; df = 5,150; *P* = 0.002), but log-transformed data were homoscedastic (Brown-Forsythe test, *F* = 1.77; df = 5,150; *P* = 0.12) and normally distributed (Kolmogorov-Smirnov *D* statistics; all *P* values > 0.20). Parametric ANCOVAs with the weather covariate (*F* = 5.94; df = 5,149; *P* < 0.0001) and the temperature covariate (*F* = 7.56; df = 5,149; *P* < 0.00001) were significant. Four pairwise comparisons were significant: Incandescent > Warm LED, Incandescent > Cool LED, Incandescent > Yellow, and CFL > Yellow.

The rate of Lepidoptera (L.) captured was different between the lamp treatment groups (total n = 901 captured). The mean ± SE nightly captures/hr was 1.16 ± 0.18 for Incandescent, 0.65 ± 0.11 for CFL, 1.02 ± 0.15 for Halogen, 0.26 ± 0.06 for Warm LED, 0.32 ± 0.07 for Cool LED, and 0.16 ± 0.03 for Yellow. Sample distributions were heteroscedastic (Brown-Forsythe test, *F* = 9.82; df = 5,150; *P* < 0.00001), as were the log-transformed data (Brown-Forsythe test, *F* = 6.91; df = 5,150; *P* < 0.00001). A nonparametric ANCOVA with the weather covariate was significant (Quade’s extension of the Kruskal-Wallis *H* = 53.29; *P* < 0.00001). Pairwise comparisons were performed with a Bonferroni corrected α level = 0.05/15 = 0.0033. Seven comparisons were significant (Figure 3).

**Fig. 3.**
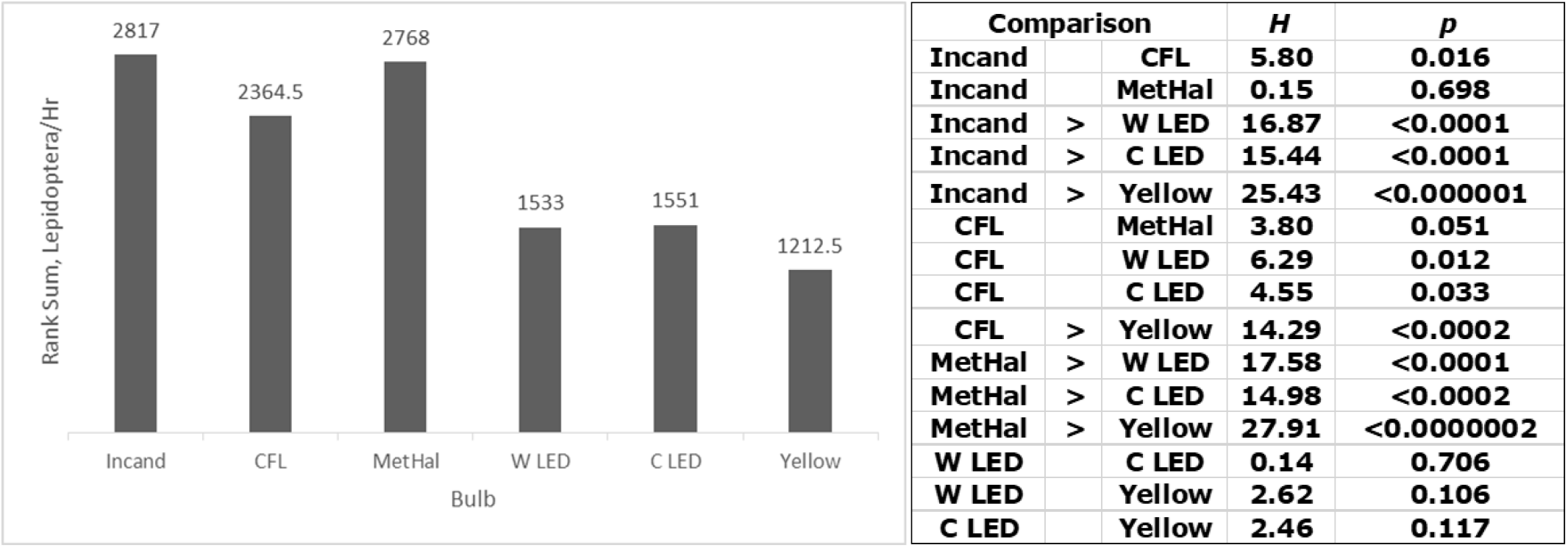
Lepidoptera/hour rank sum differences and pairwise comparisons. α = 0.05/15 = 0.0033.

The rate of Hemiptera (L.) captured was different between the lamp treatment groups (total n = 879 captured). The mean ± SE nightly captures/hr was 0.76 ± 0.13 for Incandescent, 0.68 ± 0.14 for CFL, 0.44 ± 0.10 for Halogen, 0.35 ± 0.08 for Warm LED, 0.46 ± 0.09 for Cool LED, and 0.74 ± 0.13 for Yellow. One sample distribution (Cool LED) showed evidence of nonnormality (Kolmogorov-Smirnov *D* = 0.27; 0.02 < *P* < 0.05). Log-transformed data were homoscedastic (Brown-Forsythe test, *F* = 1.40; df = 5,150; *P* = 0.23) and normally distributed (Kolmogorov-Smirnov *D* statistics; all *P* values > 0.20 except one 0.20 > *P* > 0.10). A parametric ANOVA on the log-transformed data yielded *F* = 2.201; df = 5,150; *P* = 0.057. The weather covariate actually reduced the value of *F*, to 1.63 (*P* = 0.16) because this covariate lacked homogeneity of regression slopes with this dependent variable. The temperature covariate, however, yielded *F* = 4.167; df = 5,149; *P* < 0.002. Two pairwise comparisons were significant: Incandescent > Warm LED and Yellow > Warm LED.

The rate of Dermaptera (Leach) captured was different between the lamp treatment groups (total n = 195 captured). The mean ± SE nightly Dermaptera captures/hr was 0.01 ± 0.006 for Incandescent, 0.09 ± 0.031 for CFL, 0.03 ± 0.015 for Halogen, 0.07 ± 0.030 for Warm LED, 0.06 ± 0.022 for Cool LED, and 0.48 ± 0.113 for Yellow (see Figure 4). Rank-based statistics were used here because all bulbs except Yellow had many zero scores. A nonparametric ANCOVA was significant (*H* = 21.66; *P* < 0.001). Two pairwise comparisons were significant: Yellow > Incandescent (*H* = 17.76; *P* < 0.0001) and Yellow > Halogen (*H* = 11.04; *P* < 0.0001). The other three comparisons with Yellow were noteworthy: 1) vs. CFL, *H* = 8.19; *P* = 0.0042; 2) vs. Warm LED, *H* = 5.18; *P* = 0.0228; and 3) vs Cool LED, *H* = 7.83; *P* = 0.005. Further analyses of Dermaptera are presented in Table 3.

**Fig. 4.**
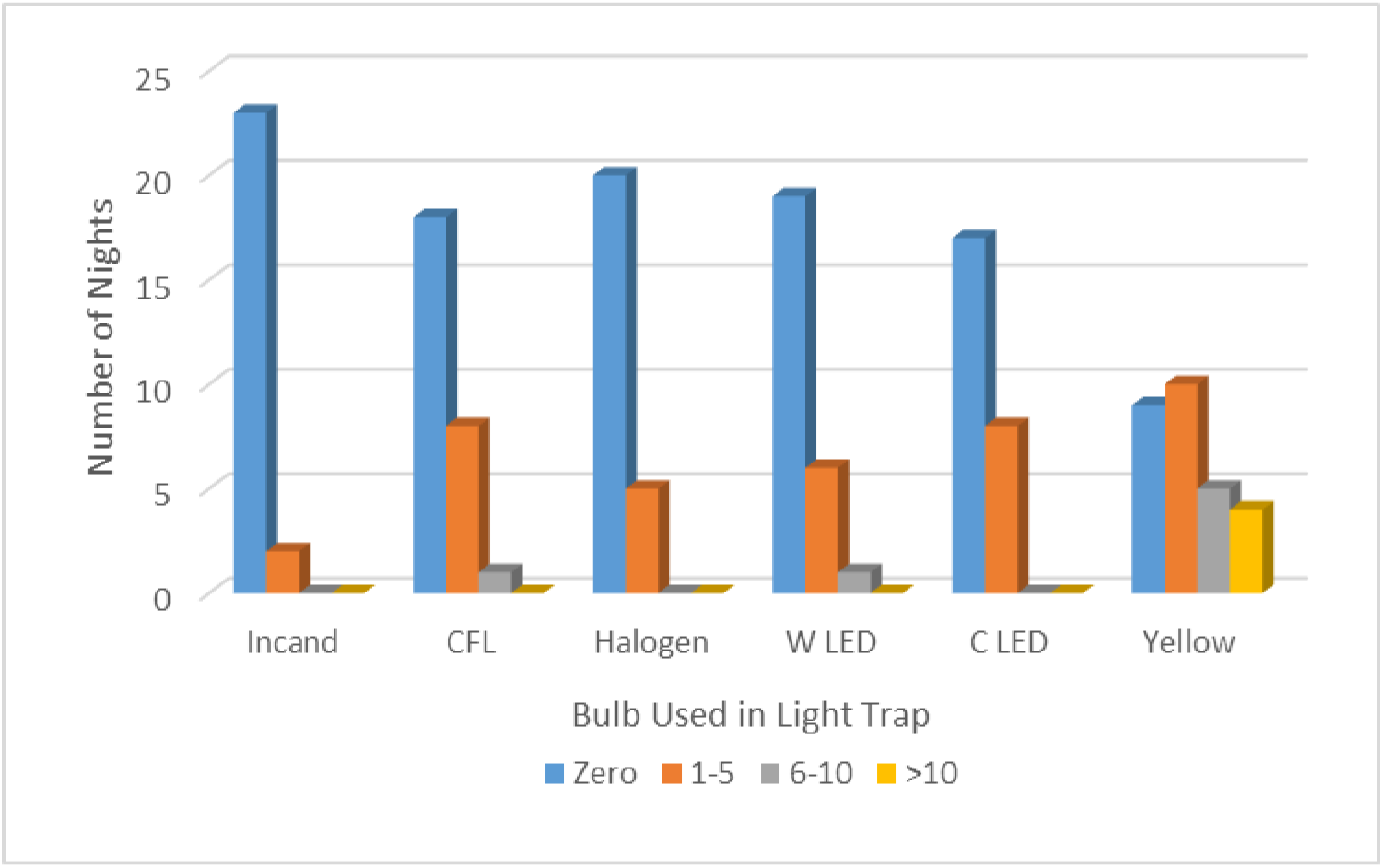
The Yellow “Bug” Light attracted far greater numbers of Dermaptera than all of the other bulbs. The color key at the bottom indicates the number of Dermaptera captured during one night of trapping.

**Table 3.**
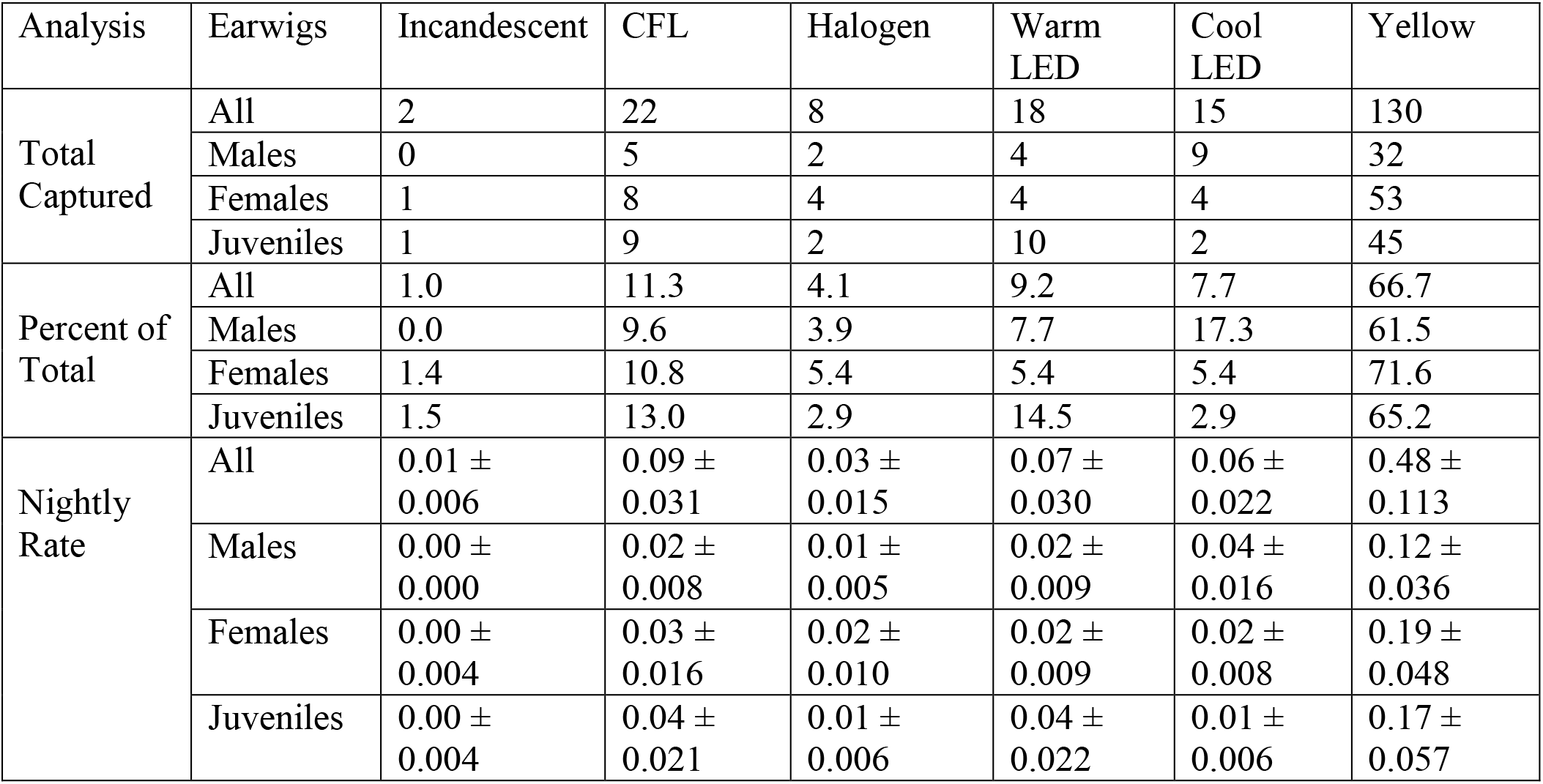
Dermaptera captures by bulb type, age, and gender. “Nightly Rate” is the mean captures per hour of darkness across the nights a given bulb was used, ± SE.

The rate of Hymenoptera (L.) captured was not different between the lamp treatment groups (total n = 2669 captured). The mean ± SE nightly captures/hr was 2.16 ± 0.54 for Incandescent, 2.11 ± 0.41 for CFL, 1.70 ± 0.33 for Halogen, 1.83 ± 0.36 for Warm LED, 1.51 ± 0.23 for Cool LED, and 1.26 ± 0.18 for Yellow. Sample distributions were homoscedastic (Brown-Forsythe test, *F* = 1.34; df = 5,150; *P* = 0.25) and normally distributed (Kolmogorov-Smirnov *D* statistics; all *P* values > 0.20 except one 0.20 > *P* > 0.10). A parametric ANCOVA with the weather covariate was not significant (*F* = 1.04; df = 5,149; *P* = 0.40), as was an ANCOVA with the temperature covariate (*F* = 1.93; df = 5,149; *P* = 0.09).

The rate of Coleoptera (L.) captured was not different between the lamp treatment groups (total n = 1484 captured). The mean ± SE nightly captures/hr was 1.33 ± 0.30 for Incandescent, 1.11 ± 0.17 for CFL, 0.89 ± 0.15 for Halogen, 0.59 ± 0.11 for Warm LED, 1.10 ± 0.23 for Cool LED, and 0.83 ± 0.19 for Yellow. Sample distributions were homoscedastic (Brown-Forsythe test, *F* = 1.74; df = 5,150; *P* = 0.13) and normally distributed (Kolmogorov-Smirnov *D* statistics, all *P* values > 0.10 except one 0.10 > *P* > 0.05). A parametric ANCOVA with the weather covariate was not significant (*F* = 1.50; df = 5,149; *P* = 0.194), as was an ANCOVA with the temperature covariate (*F* = 2.13; df = 5,149; *P* = 0.065).

## Discussion

This study directly compared insect attraction to all six of the major types of low-wattage light bulb used for lighting small areas around residential exteriors. In agreement with previous research (Justice & Justice 2016; Eisenbeis & Eick 2011), the data suggest that a large-scale switch to LED bulbs, especially those with a warm color temperature, could greatly diminish the effects of night lighting on insect behavior and mortality. Further, the use of yellow “bug” lights, in contrast to their marketing, could attract earwigs and other minor pests. Each type of bulb already has a constellation of environmental concerns, alongside which relative insect attraction can now be considered.

Incandescent bulbs are being phased out of retail markets due to their energy inefficiency. A widespread reduction in their use will likely benefit insects, as incandescent bulbs produced the highest mean capture rate for all orders combined as well as all individual orders examined. Mean differences between incandescent and at least one other bulb were statistically significant in four cases: total insects and spiders, Diptera, Lepidoptera, and Hemiptera. These groups contain a number of major and minor pests, vectors, and nuisances. The hot operating temperature of incandescent bulbs means that, in addition to attracting insects away from their normal behaviors, desiccation and burning are probably more likely around them. Other bulbs, in contrast, could possibly spare more insects and allow them to leave and resume normal activities. It is thus distinctly possible that incandescent bulbs could be phased out of common household use, largely benefitting insects, yet used strategically in limited circumstances to attract and destroy pests.

Halogen bulbs are technologically very similar to incandescent. A halogen-laden gas inside a quartz housing allows for hotter, more optimal operating temperatures, longer life, a slight increase in energy efficiency, and a slight increase in color temperature. The lamp used in the present study was encapsulated in a bulb-shaped glass housing for cosmetic and convenience purposes. This allowed for a more equitable comparison with the other bulb-shaped lamp housings in this study, and also prevented insects from directly contacting the extremely hot lamp itself. Given the similarities between them, it is not surprising that the insect capture rate of the halogen bulb was never significantly different from that of the incandescent bulb. Significant mean differences suggest halogen bulbs attract significantly more Lepidoptera than the LED bulbs and the yellow “bug” light. Given, in addition, the marginal differences from the incandescent bulbs, their use of halogen gases, and their dangerously hot temperatures, there is not much to recommend the use of halogen bulbs from an ecological or consumer standpoint.

CFLs, in all analyses, had middle to high mean capture rates, and these were never significantly less than incandescent or halogen. CFLs also attracted significantly more total insects and spiders than the warm color temperature LED bulb, and more Diptera and Lepidoptera than the yellow “bug” light. They are somewhat more energy efficient than incandescent and halogen bulbs, but this environmental benefit could be countered by improper disposal, which can release their mercury gas.

Yellow “Bug” Lights are usually marketed as lights that will not attract insects, and sometimes even as lights that repel insects. In the present study, the yellow “bug” light produced the second-lowest mean capture rate of total insects, and this was significantly lower than the incandescent bulb. Thus, installation of these bulbs on porches etc. will likely reduce the numbers of insects attracted to these areas, especially if they replace an incandescent bulb. However, there are three major caveats regarding their use. First, the light they produce is generally not regarded as pleasant to the people using them. As with sodium streetlights, bathing people and objects in yellow light can distort appearances. “Whiter” light with a broader mix of wavelengths is understandably preferred and occasionally necessary. Second, the yellow light is not produced by a fundamentally different technology. “Bug” lights are often simply incandescent bulbs or CFLs within a housing that filters out all but yellow wavelengths. Thus, similar problems such as mercury release or high operating temperatures could be present, and energy efficiencies could be even worse because usable light is being produced within but not being allowed to escape the bulb. Third, the CFL-based “bug” light used in the present study attracted more Dermaptera than the other bulbs (especially the incandescent and halogen bulbs), and significantly more Hemiptera than the warm color temperature LED. Dermaptera and many Hemipterans can be minor pests around residences. Thus, yellow “bug” lights are probably not the best choice from both ecological and consumer perspectives, but their use in traps for Dermaptera and Hemiptera should be considered. Insect traps can also destroy large numbers of beneficial species (Frick & Tallamy 1996), so exactly what the yellow “bug” lights attract should be assessed down to at least the Family level.

LEDs are the most recent bulbs to be made widely available to consumers, and appear to be an improvement upon previous technologies from both a consumer and ecological point of view. The cost of LEDs is currently relatively high, but their energy efficiency and lifespans are favorable compared to other bulbs. Features such as instant-on and dimming, which are attractive to consumers, are now available with LEDs. Also, their technology creates lighting that is strongly directional; this allows for more precise lighting of a given space and makes it easier to reduce the unwanted zenith-directed light that contributes to skyglow. Warm LEDs have already been shown to be insect-friendly in the higher wattages of street lights (Eisenbeis & Eick 2011). Among the lower-wattage residential-style bulbs used in the present study and in Justice & Justice (2016), the warm and cool color temperature LEDs had the lowest mean capture rates of total insects and spiders among the bulbs producing “white” light (i.e. excluding the yellow “bug” light). Their mean capture rates were significantly lower than at least one other bulb in the analyses comparing total insects, Diptera, Lepidoptera, and Hemiptera. Although the mean capture rates of these two bulbs were never significantly different from each other, the warm color temperature LED should be favored over the cool color temperature LED for residential use. Blue wavelengths tend to contribute more to light pollution, and these are attenuated to create the warm color temperature.

One caveat is that widespread adoption of a product magnifies its impacts through all stages of its life cycle. Before any recommendations are made to consumers, life cycle assessments and moral purchasing analyses of LED bulbs should pay careful attention to the extraction and disposition of the minerals needed for their phosphors and solid-state electronics. Of course, fewer considerations of this sort would be needed if stray photons were more widely regarded as municipal waste and potential pollutants.

## Acknowledgments

Bascom Deaver and Acar Işin, Department of Physics, University of Virginia, Charlottesville, Virginia, USA provided invaluable assistance, expertise, and equipment for the spectrophotometry. The suggestion to use the Yellow “Bug” Light came from Mark Deyrup, Archbold Biological Research Station, Lake Placid, FL. During the course of this research, Michael J. Justice was supported by a travel grant from the Virginia Department of Education.

## Notes

### Competing Interest Statement

The authors have declared no competing interest.

